# Relative stability of mRNA and protein severely limits inference of gene networks from single-cell mRNA measurements

**DOI:** 10.1101/2022.03.31.486623

**Authors:** Tarun Mahajan, Michael Saint-Antoine, Roy D. Dar, Abhyudai Singh

## Abstract

Inference of gene regulatory networks from single-cell expression data, such as single-cell RNA sequencing, is a popular problem in computational biology. Despite diverse methods spanning information theory, machine learning, and statistics, it is unsolved. This shortcoming can be attributed to measurement errors, lack of perturbation data, or difficulty in causal inference. Yet, it is not known if kinetic properties of gene expression also cause an issue. We show how the relative stability of mRNA and protein hampers inference. Available inference methods perform benchmarking on synthetic data lacking protein species, which is biologically incorrect. We use a simple model of gene expression, incorporating both mRNA and protein, to show that a more stable protein than mRNA can cause loss in correlation between the mRNA of a transcription factor and its target gene. This can also happen when mRNA and protein are on the same timescale. The relative difference in timescales affects true interactions more strongly than false positives, which may not be suppressed. Besides correlation, we find that information-theoretic nonlinear measures are also prone to this problem. Finally, we demonstrate these principles in real single-cell RNA sequencing data for over 1700 yeast genes.

## I. Introduction

According to the “central dogma” of molecular biology [1], genes on DNA are transcribed into messenger RNA (mRNA), which are translated into proteins. The proteins carry out various functions within the cell including regulation of expression of other genes. These regulatory relationships form gene regulatory networks (GRNs), which control the complexity of cellular life [2], [3], and malfunctions in GRNs can lead to diseases like cancer [4].

Understanding GRN function and structure is essential for cell biologists, and the inference of their topology from static transcriptomic data is important [5]. Researchers have used statistical relationships between levels of mRNA to identify the underlying GRN. These statistical methods include correlation [6], regression [7]–[9], information-theoretic techniques [10]–[14], Bayesian techniques [15], [16], and more [17]–[23]. Benchmarking studies have assessed the comparative performance of different methods [24]. An excellent review of GRN inference methods can be found in [25].

Many GRN inference techniques make the assumption that mRNA and protein counts for a given gene are correlated, and use mRNA transcript abundance as a proxy for protein abundance, which is difficult to measure in a highthroughput manner. However, in experiments the correlation between mRNA and corresponding protein counts can be weak [26]. In this paper, we use a model of gene expression to explore the impact of relative stability of mRNA and protein on the correlation between their abundances. We also explore information-theoretic measures–mutual information (MI) [27] and the phi-mixing coefficient [28]–using stochastic simulations [29]. MI quantifies the uncertainty in individual and joint distributions of random variables [27]. The phi-mixing coefficient is a measure of the statistical dependence of two random variables based on the difference between their conditional and unconditional probability distributions [28]. Both MI ([10]–[14]) and the phi-mixing coefficient ([30]) have been used for GRN inference. Finally, we demonstrate the established principles on a real singlecell RNA sequencing (scRNA-seq) dataset for yeast, *S. cerevisiae*.

We show that when protein is more stable than mRNA, GRN cannot be reliably inferred at the single-cell mRNA level. Even when the true GRN edges are identifiable, the false positive edges will dominate in correlation values. Collectively, this establishes a protein-mRNA lifetime-dependent loss in GRN signature in single-cell data.

## II. A simple model of gene expression

We start with a simple model of gene expression. mRNA *M* is produced and degraded via a 1-dimensional Poisson birth-death process. Uppercase and lowercase letters represent a species and its molecular count, respectively; *M* and *m* are the mRNA and its count, respectively. Production of *M* is a Poisson birth process with rate *k*_*m*_*F*(*β*), and each event creates mRNA in bursts of size *β*, which is distributed according to any arbitrary positive-valued probability distribution *F*(*β*). Protein *P* is created and degraded in a 1-dimensional conditional Poisson birth-death process. Each *M* molecule is translated into a molecule of *P* via a conditional Poisson birth process at a rate *k*_*p*_. *M* and *P* are degraded as Poisson processes at rates 1*/τ*_*m*_ and 1*/τ*_*p*_, respectively. *τ*_*m*_ and *τ*_*p*_ are the respective average lifetimes of *M* and *P*. For the rest of the paper, we assume that *τ*_*m*_ is constant, and *τ*_*p*_ = *τ τ*_*m*_. This allows us to change *τ*_*p*_*/τ*_*m*_ by varying only *τ*. The chemical reaction network for this system is

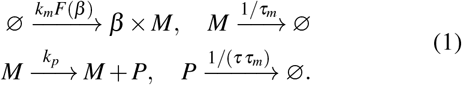

The time evolution of the joint probability distribution for *m* and *p* in (1) is given by the following chemical master equation (CME) [31]:

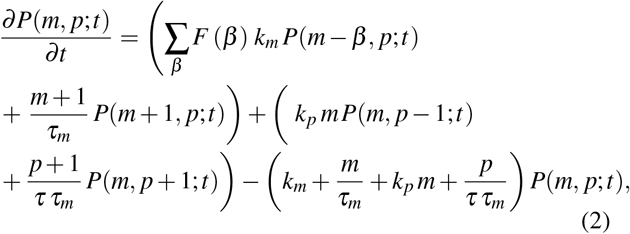

where *P*(*m, p*; *t*) is the joint probability distribution for *m* and *p* at time *t*. (2) can be solved exactly for the steady-state moments [31]–[33]. The first order moments are given by

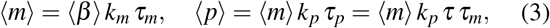

where the angle brackets represent statistical expectation. The second order moments involving only one species can be written as

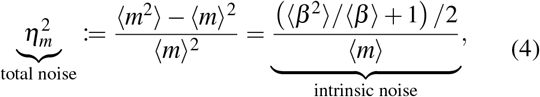

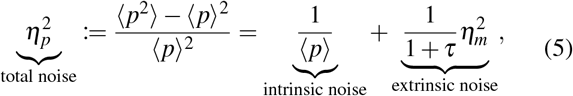

where 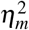 and 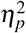 are the total noise or expression fluctuation, for *M* and *P*, respectively [32], [33]. Intrinsic noise is the fluctuation in *m* or *p* caused by the discrete birth-death events for *M* or *P*, respectively. Intrinsic noise for any mRNA species is given by (4) [32], [33]. Intrinsic noise for any protein species is given by the first term on the right-hand size (RHS) in (5) [32], [33]. Extrinsic noise in (5) is the noise propagated from *m* to *p*. The decomposition of total noise into intrinsic and extrinsic in (5) is fairly standard in noisy expression research [32]–[36]. The numerator for intrinsic noise in (4),

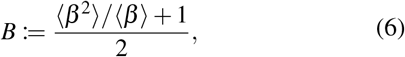

is the average contribution to noise from birth and death events for *M* [34], [35]. We propose that

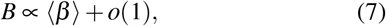

where *o* is the Little-o notation. If *β* is deterministic, *B* = (⟨*β*⟩ + 1) */*2. If *β* is distributed geometrically, as is known for many genes in different species [37], [38], *B* = ⟨*β*⟩. *B* is an estimate of mean-independent intrinsic noise for mRNA, and we modulate noise by varying *B*. We vary *B* by changing ⟨*β*⟩ ((7)). Whenever ⟨*β*⟩ is increased/decreased by a factor, we decrease/increase *k*_*m*_ by the same factor to keep first order moments constant at fixed *τ*.

We are interested in the dependence of steady-state species correlations on *B*, and *τ*. Since the moments in (3)- (5) are dependent on *B* and *τ*, we obtain the expression for correlation between *m* and *p*, cor(*m, p*), as a function of these moments.

### Proposition 1 (mRNA-protein correlation in a single gene)

*For a single gene* (1), *steady-state correlation between m and p is*

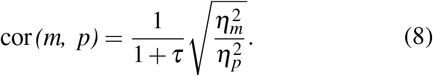

(8) is obtained by solving (2) [31]–[33]. We examine the behavior of cor(*m, p*) when *τ* is fixed, and *B* is variable, and establish the following upper bound:

### Theorem 2 (Upper bound on mRNA-protein correlation in a single gene)

*For a single gene* (1), *steady-state correlation between m and p is bounded from above by*

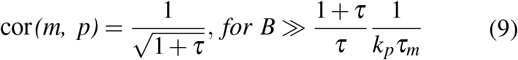

cor(*m, p*) is an increasing function of 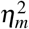, which is an increasing function of *B* (see (4) and (6)). Consequently, cor(*m, p*) is an increasing function of *B*, and reaches its maximum value in (9) when the second term on the right hand side (RHS) in (5) is much larger than the first term. This gives us the constraint on *B* in (9) while using (3)-(4).(9) is depicted by the green curve in Fig. 1b. There is a *τ*-mediated loss in correlation as protein becomes more stable than mRNA. For extremely stable proteins, cor(*m, p*) might completely vanish. Actual cor(*m, p*) values will always be lower than (9). The gap between the upper bound and the true cor(*m, p*) is governed by the relative amounts of intrinsic and extrinsic noises in *p*. This is evident from (8) and (9). cor(*m, p*) reaches the upper bound only when *B* is large enough. However, if *B* is low or moderate, cor(*m, p*) can deviate significantly. cor(*m, p*)’s dependence on *τ* and *B* is shown in Fig. 1b. Further, we have also validated the analytical result using exact stochastic simulations in Fig. 1c. Dependence of cor(*m, p*) on *τ* and *B* motivates the central theme of the paper. We next explore whether this dependence propagates to downstream genes in small GRN topologies.

**Fig. 1:**
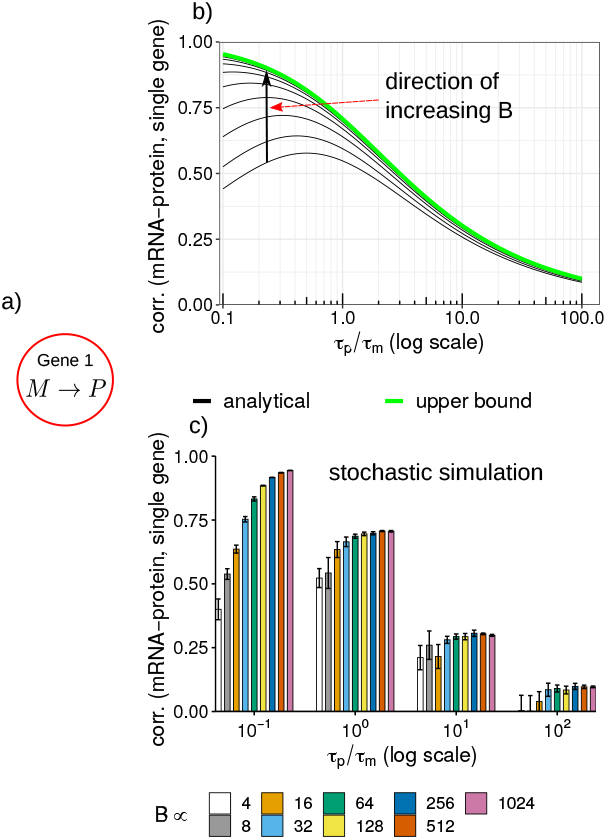
**Plot of mRNA-protein correlation, cor(*m, p*)**, for a **single gene** in (1) as a function of *B*, and *τ*. (a) GRN being considered–a single gene with its mRNA, *M*, and protein, *P*. (b) Analytical curves (black curves) obtained from (8) (one curve per *B* value). Nine different values were used for *B*: *B* ∝ ⟨*β*⟩ ∈ 4 × {2^0^, 2^1^, …, 2^8^}. The upper bound in (9) is depicted by the green curve. (c) cor(*m, p*) as a function of *B* and *τ* using exact stochastic simulations [29]. *B* was varied over the same range as before. The other kinetic parameters are: *k*_*m*_ = 0.0282, *τ*_*m*_ = 1*/*0.0025, *k*_*p*_ = 1.2*/τ*_*m*_. *k*_*m*_ and *τ*_*m*_ are for the nanog gene from mouse embryonic stem cells ([39]). We assume that *β* is deterministic. Error bars show one standard deviation.

## III. Loss of Correlation for Simple GRNS

### Count correlations in a two-gene cascade

For a two-gene cascade (Fig. (2)a), gene 1 has mRNA *M*_1_ and protein *P*_1_, and gene 2 has mRNA *M*_2_ and protein *P*_2_. *P*_1_ regulates the production of *M*_2_. All kinetic parameters are identical for the two genes. *M*_1_ is created and degraded via a 1-dimensional Poisson birth-death process. All the other species are created and degraded via 1-dimensional conditional Poisson birth-death processes. The chemical reaction network is

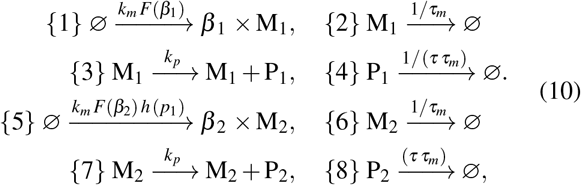

where *β*_1_ and *β*_2_ are the respective burst sizes for *M*_1_ and *M*_2_, *P*_1_ regulates *M*_2_ via *h* (*p*_1_), and numbers (in curly braces) represent an ordering on the reactions. The CME for (10) in compact form is [31]

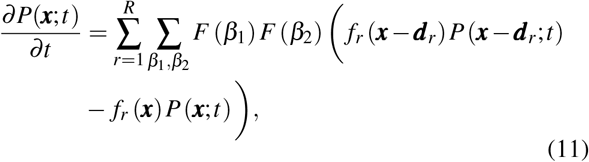

where *R* = 8 is the total number of reactions in (10), ***x*** = (*m*_1_, *p*_1_, *m*_2_, *p*_2_)^*T*^, *f*_*r*_ (***x***) is the propensity for reaction 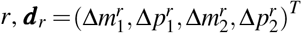 is a vector containing the jump in species count because of reaction *r*, and *P*(***x***; *t*) is the joint probability distribution for ***x***. *β*_1_ and *β*_2_ independent. We define

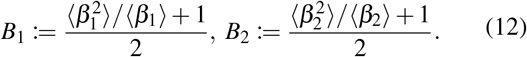

as the average contributions to noise from birth and death processes for *M*_1_ and *M*_2_, respectively ([32]–[35]):

(11) can be solved exactly for steady-state moments if *h* (*p*_1_) is a linear function [31]–[33]. For nonlinear *h* (*p*_1_), we solve (11) approximately using the linear noise approximation (LNA) approach [31]–[35], [40], [41]. For LNA, we linearize *h*(*p*_1_) around a deterministic concentration [32], [33], [41]. Then, evolution of the first-order moments is obtained using the extended moment generator from [42]:

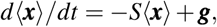

where *S* is a diagonal matrix of inverse lifetimes, and **g** is the vector of species production rates. Time evolution of the mean-normalized covariance matrix Σ, with entries Σ_*i j*_ = (⟨*x*_*i*_*x* _*j*_⟩ − ⟨*x*_*i*_⟩ ⟨ *x* _*j*_⟩)*/*(⟨*x*_*i*_⟩ ⟨*x* _*j*_ ⟩) is given by ([31], [34], [35], [40], [41])

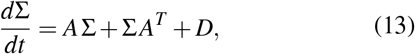

where *A* and *D* are the mean-normalized jacobian and diffusion matrices, respectively. Ordering for the rows and columns of *S, A, D* and Σ corresponds to the species order in ***x***. The non-zero entries of *A* encode the structure of the expanded GRN including both mRNA and protein; there exists a regulation edge from species *j* to *i* iff *A*_*i j*_ ≠ 0 ([31]–[35], [40], [41]). At steady state, first-order moments are given by

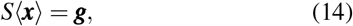

and the Σ is given by the following Lyapunov equation ([31]–[35]):

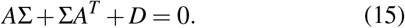

**Fig. 2:**
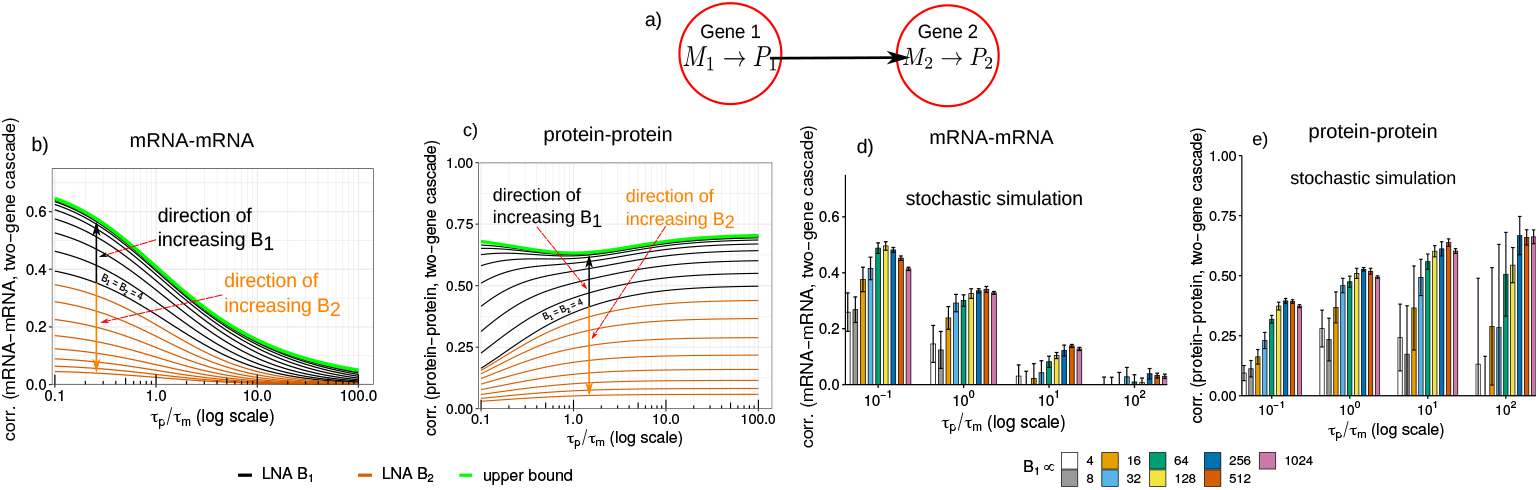
**Plot of mRNA-mRNA, cor(*m*_1_, *m*_2_), and protein-protein, cor(*p***_**1**_, ***p***_**2**_**), correlation**, for a **two-gene cascade** in (10) as a function of *B*_1_, *B*_2_, and *τ*. (a) GRN being considered–a two gene cascade. Analytical curves (black (one curve per value of *B*_1_, while *B*_2_ ∝⟨ *β*_2_⟩ = 4) and yellow (one curve per value of *B*_2_, while *B*_1_ ∝ ⟨*β*_1_⟩ = 4) curves) for cor(*m*_1_, *m*_2_) and cor(*p*_1_, *p*_2_) are shown in (b) and (c), respectively. The upper bounds in (17) and (22) are depicted by the green curves in (b) and (c), respectively. cor(*m*_1_, *m*_2_) and cor(*p*_1_, *p*_2_) as functions of *B*_1_ and *τ* using exact stochastic simulations [29] are shown in (d) and (e), respectively. *B*_1_ was varied while *B*_2_ was held constant (both in the same ranges as before). The values of other kinetic parameters are the same as mentioned in the caption for Fig. 1. Further, we assume that 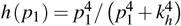, where *k*_*h*_ was selected such that at steady state *h* (⟨*p*_1_ ⟩) = 0.5 for all values of the kinetic parameters. Error bars show one standard deviation.

(15) gives all the second order moments ([31]–[35], [41]). We are interested in the dependence of steady-state correlation between *m*_1_ and *m*_2_, cor(*m*_1_, *m*_2_), on *B*_1_, *B*_2_ and *τ*.

#### Proposition 3 (mRNA-mRNA correlation in a two-gene cascade)

*For the two-gene cascade in* (10), *correlation between m*_1_ *and m*_2_ *is*

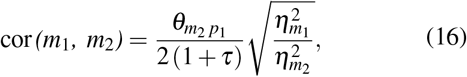

where 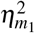 and 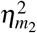 are the total noises in *m*_1_ and *m*_2_, respectively, which are obtained from (15) ([31]–[35], [41]). 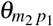 is the log-sensitivity of *m*_2_ to changes in *p*_1_ at steady state ([32]–[35]). Assume that *h* (*p*_1_) is saturating and *h* (⟨*p*_1_⟩) is independent of ⟨*p*_1_⟩. Then, 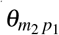 is also independent of ⟨*p*_1_⟩ at steady-state [32]–[35]. (16) is obtained by solving (15) ([31]–[33], [41]). We next establish the following upper bound on cor(*m*_1_, *m*_2_):

#### Theorem 4 (Upper bound on mRNA-mRNA correlation in a two-gene cascade)

*For the two-gene cascade in* (10), *correlation between m*_1_ *and m*_2_ *is bounded from above by*

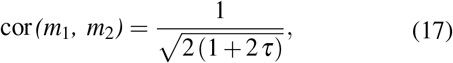

*when*

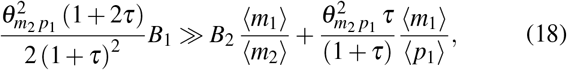

From (15), we obtain ([32], [33]),

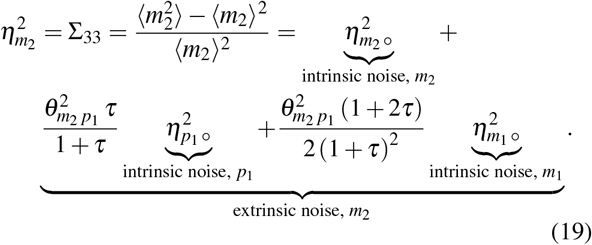

From (16), (19) and 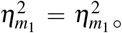, cor(*m*_1_, *m*_2_) is an increasing function of 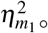, and achieves the upper bound in (17) when the third term on the RHS in (19) is much larger than the first two terms. (17) is depicted by the green curve in Fig. 2b. There is a *τ*-mediated loss in cor(*m*_1_, *m*_2_). As protein becomes more stable than mRNA, cor(*m*_1_, *m*_2_) decays much faster than cor(*m*_1_, *p*_1_). For extremely stable proteins, cor(*m*_1_, *m*_2_) will completely vanish. Actual cor(*m*_1_, *p*_1_) could be much less than (17) (Fig. 2).

(18) defines a tug-of-war between *B*_1_ and *B*_2_. Their relative magintudes dictate the gap between (17) and the actual cor(*m*_1_, *m*_2_). If *B*_1_ is much higher than *B*_2_, (18), then cor(*m*_1_, *m*_2_) will reach its upper bound (17) (see the black curves in Fig. 2b). However, if *B*_1_ is much smaller than *B*_2_, cor(*m*_1_, *m*_2_) can vanish even when mRNA is more stable than protein, and *τ <* 1 (check the yellow curves in Fig. 2b). We also performed exact stochastic simulations ([29]) to verify (16) and (17), Fig. 2d. Next, we examine the steady-state correlation between *p*_1_ and *p*_2_.

#### Proposition 5 (protein-protein correlation in a two-gene cascade)

*For the two-gene cascade* (10), *correlation between p*_1_ *and p*_2_ *is*

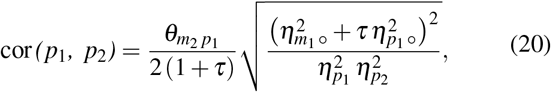

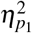 is obtained from (5) by replacing 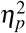 and 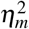 with 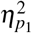 and 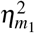, respectively, and from (15) ([32], [33])

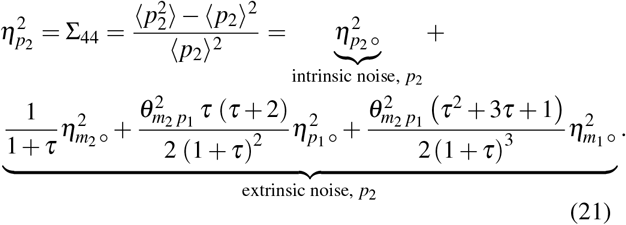

(20) is obtained from (15) [31]–[33]. cor(*p*_1_, *p*_2_) is an increasing function of 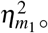. Consequently, cor(*p*_1_, *p*_2_) is an increasing function of *B*_1_, which establishes the following upper bound on cor(*p*_1_, *p*_2_):

#### Theorem 6 (Upper bound on protein-protein correlation in a two-gene cascade)

*For the two-gene cascade* (10), *correlation between p*_1_ *and p*_2_ *is bounded from above by*

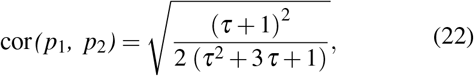

*when*

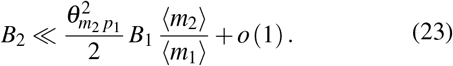

Theorem 6 can be proved by substituting intrinsic noise terms in (20) to show that the upper bound is reached when the fourth term in the RHS of (21) is much larger than the first three terms. When *B*_1_ is large to satisfy (23), cor(*p*_1_, *p*_2_) will reach its upper bound in (22) (see the black curves in Fig. 2c). For all values of *τ*, cor(*p*_1_, *p*_2_) is greater than or equal to 80% of its maximum of 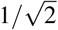 (green curve in Fig. 2c). This is in contadiction to (17) where the upper bound was a monotonically decreasing function of *τ*. The upper bound for cor(*p*_1_, *p*_2_) does not exhibit a *τ*-mediated loss in correlation. cor(*p*_1_, *p*_2_) also exhibits a tug-of-war between *B*_1_ and *B*_2_ (yellow curves in Fig. 2c). This is evident from (23) and the fact that (20) is a decreasing function of *B*_2_.

We also performed stochastic simulations [29] to demonstrate the dependence of cor(*p*_1_, *p*_2_) on *B*_1_, and *τ* (Fig. 2e).

### Count correlations in a three-gene fanout motif

Next, we show that the false positive correlation between two genes having a common upstream TF, but not regulating each other, is less susceptible to a *τ*-mediated loss of correlation. This false positive correlation is most of the time greater than the true positive correlation in a two-gene cascade. For this, we study a fanout network, which has three genes. Genes 1 and 2 satisfy (10). Gene 3 satistifes the following additional reaction channels:

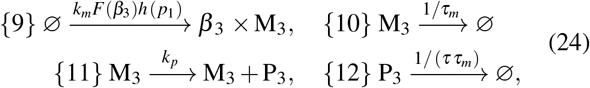

where reaction numbering has been continued from (24), *M*_3_ and *P*_3_ are the mRNA and protein for gene 3, respectively, *β*_3_ is the burst size for *m*_3_. We assume that *β*_2_ and *β*_3_ are identically distributed. We also define

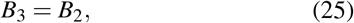

where *B*_2_ is given in (12), as the average contribution of the birth and death events in (24) to noise in *m*_3_. Genes 2 and 3 are identical, and do not regulate each other. They have a common regulator, *P*_1_. Correlation between the species of genes 2 and 3 defines false positive correlation (cor(*m*_2_, *m*_3_) and cor(*p*_2_, *p*_3_)). Now, ***x*** = (*m*_1_, *p*_1_, *m*_2_, *p*_2_, *m*_3_, *p*_3_)^*T*^. All moments upto the second-order can be obtained by solving (14) and (15) with additional species, *M*_3_ and *P*_3_. All moments for *m*_3_ amd *p*_3_ can be obtained from the moments of *m*_2_ and *p*_2_, respectively. Consequently, 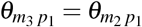. Next, we find cor(*m*_2_, *m*_3_).

#### Proposition 7 (mRNA-mRNA correlation in a three-gene fanout)

*For the three-gene fanout* (10) *and* (24), *correlation between m*_2_ *and m*_3_ *is given by*

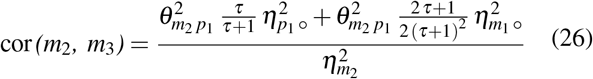

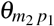 and 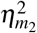 in (26) are interchangeable with 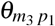 and 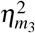, respectively. Proposition 7 is proved by solving (15) [32], [33]. On substituting (19) in (26), we see that cor(*m*_2_, *m*_3_) is an increasing function of *B*_1_, and establish the following upper bound on cor(*m*_2_, *m*_3_):

#### Theorem 8 (Upper bound on mRNA-mRNA correlation in a three-gene fanout)

*For the three-gene fanout* (10) *and* (24), *correlation between m*_2_ *and m*_3_ *is bounded from above by*

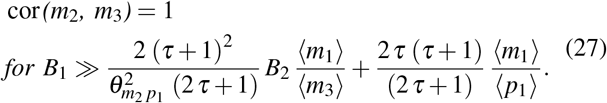

cor(*m*_2_, *m*_3_) depends on *B*_1_ through 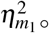. For 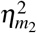 in the denominator in (26), substituting (19), we get (27). Interestingly, the upper bound (27) is independent of *τ*. This is in contrast to cor(*m*_1_, *m*_2_), where (17) which decays with *τ*. This implies that cor(*m*_2_, *m*_3_) will always dominate cor(*m*_1_, *m*_2_) and cor(*m*_1_, *m*_3_) when protein is more stable than mRNA, and can even dominate when mRNA is more stable (Fig. (3)b). Next, we directly compare cor(*m*_1_, *m*_2_) to cor(*m*_2_, *m*_3_).

**Fig. 3:**
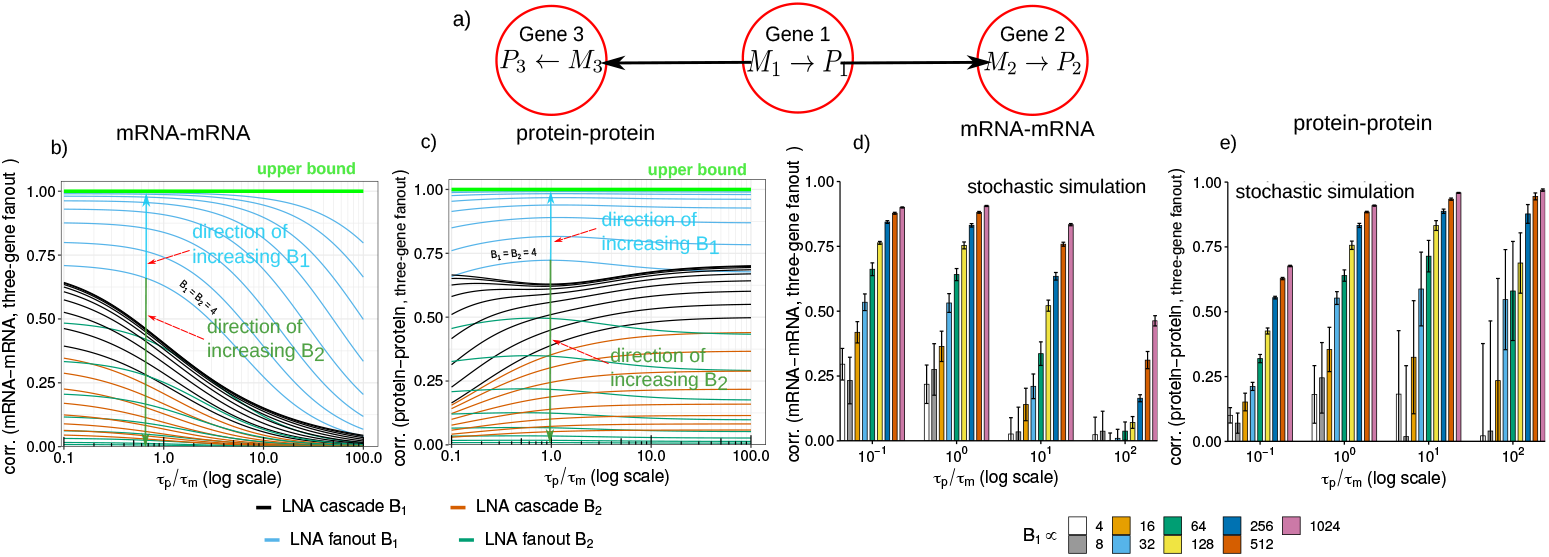
**Plot of mRNA-mRNA, cor(*m*_2_, *m*_3_), and protein-protein, cor(*p***_**2**_, ***p***_**3**_**), correlation**, for a **three-gene fanout** in (10) and (24) jointly as a function of *B*_1_, *B*_2_, and *τ*. (a) GRN being considered–a three-gene fanout Analytical curves (cyan (one curve per value of *B*_1_, while *B*_2_ ∝ ⟨*β*_2_ ⟩= 4) and dark-green (one curve per value of *B*_2_, while *B*_1_ ∝ ⟨*β*_1_ ⟩ = 4) curves) for cor(*m*_2_, *m*_3_) and cor(*p*_2_, *p*_3_) are shown in (b) and (c), respectively. The analytical curves from Figs. 2a and 2b are also shown in black and yellow curves in (b) and (c), respectively, for comparison. The upper bound of 1 is shown in green curves in (b) and (c). cor(*m*_2_, *m*_3_) and cor(*p*_2_, *p*_3_) as functions of *B*_1_ and *τ* using exact stochastic simulations [29] are shown in (d) and (e), respectively. *B*_1_ was varied while *B*_2_ was held constant (both in the same ranges as before). The values of other kinetic parameters are the same as mentioned in the caption for Fig. 1. The values of other kinetic parameters are the same as before (see caption of Fig. 1). For details on the regulation function *h* (*p*_1_), see caption of Fig. 2. Error bars show one standard deviation.

#### Theorem 9 (mRNA-mRNA correlation in two-gene cascade vs three-gene fanout)

*For the three-gene fanout* (10) *and* (24)

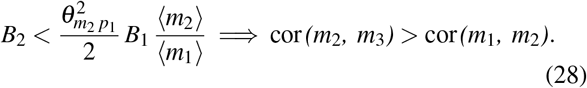

On comparing (16) and (26), while using (19), we get (28). (28) demonstrates a tug-of-war between *B*_1_ and *B*_2_, where their relative magintudes dictate whether cor(*m*_1_, *m*_2_) or cor(*m*_2_, *m*_3_) will dominate. The relative magnitudes of *B*_1_ and *B*_2_ also determine whether cor(*m*_1_, *m*_2_) and cor(*m*_1_, *m*_3_) will achieve their upper bounds (see Theorems 4 and 8). When (18) and (27) are true, cor(*m*_1_, *m*_2_) and cor(*m*_2_, *m*_3_) achieve their upper bounds. However, at the same time, cor(*m*_2_, *m*_3_) will begin to dominate cor(*m*_1_, *m*_2_). This implies that kinetic conditions which allow inference of the GRN also confound inference via the false positive edges.

The dependence of cor(*m*_2_, *m*_3_) on *B*_1_ (cyan curves), *B*_2_ (dark-green curves) and *τ* based on (26) is shown in Fig. 3b. We have also shown the respective curves for cor(*m*_1_, *m*_2_) based on (16) there for the sake of comparison. We also performed exact stochastic simulation to verify these results as shown in Figs. 3d. Next, we study the correlation between *p*_2_ and *p*_3_, cor(*p*_2_, *p*_3_).

#### Proposition 10 (protein-protein correlation in a three--gene fanout)

*For the three-gene fanout* (10) *and* (24), *correlation between p*_2_ *and p*_3_ *is*

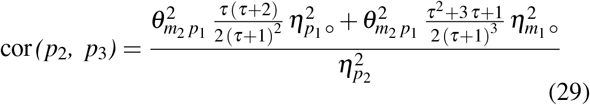

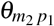 and 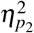 in (29) are interchangeable with 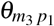 and 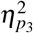, respectively. Proposition 10, is proved by solving (15) [32], [33]. Like before, we assume that *B*_1_, *B*_2_ and *B*_3_ are varied in a manner which preserves the first order moments at fixed *τ*. On substituting (21) in (29), we see that cor(*p*_2_, *p*_3_) is an increasing function of *B*_1_. Now, we establish the following upper bound on cor(*p*_2_, *p*_3_):

#### Theorem 11 (Upper bound on protein-protein correlation in a three-gene fanout)

*For the three-gene fanout* (10) *and* (24), *correlation between p*_2_ *and p*_3_ *is bounded from above by*

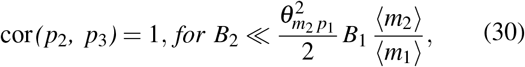

cor(*p*_2_, *p*_3_) depends on *B*_1_ through 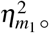. For 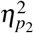 in the denominator in (29), substituting (21), we get (30) for reaching the upper bound. Similar to cor(*m*_2_, *m*_3_), the upper bound for cor(*p*_2_, *p*_3_) is independent of *τ*. Therefore, it is possible that cor(*p*_2_, *p*_3_) might dominate cor(*p*_1_, *p*_2_) and cor(*p*_1_, *p*_3_). Next, we directly compare cor(*p*_2_, *p*_3_) to cor(*p*_1_, *p*_2_):

#### Theorem 12 (Protein-protein correlation in two-gene cascade vs three-gene fanout)

*For the three-gene fanout* (10) *and* (24),

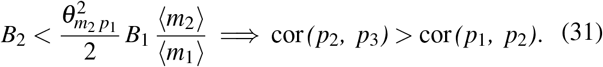

On comparing (20) and (29), while using (19), we get (31). Similar to Theorem 9, Theorem 12 demonstrates a tug-of-war between *B*_1_ and *B*_2_, where their relative magintudes dictate whether the true positive correlation cor(*p*_1_, *p*_2_) (cor(*p*_1_, *p*_3_)) or the false positive correlation cor(*p*_2_, *p*_3_) will dominate in single-cell protein measurements. Strong bursting in *M*_1_ relative to *M*_2_ can make cor(*p*_2_, *p*_3_) dominate.

The kinetic regime which allows cor(*p*_1_, *p*_2_) and cor(*p*_1_, *p*_3_) to achieve their upper bounds also allows cor(*p*_2_, *p*_3_) to reach its upper bound (compare the constraints in Theorem 6 and Theorem 11). However, the upper bound of cor(*p*_2_, *p*_3_) is larger than that of cor(*p*_1_, *p*_2_) and cor(*p*_1_, *p*_3_): 1 vs 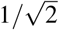. Even though protein-protein correlations are not subject to *τ*-mediated loss in correlation, yet the relative gap between the upper bounds for cor(*p*_2_, *p*_3_) and cor(*p*_1_, *p*_2_) or cor(*p*_1_, *p*_3_) will cause the false positive correlation cor(*p*_2_, *p*_3_) to dominate the true positive correlations cor(*p*_1_, *p*_2_) and cor(*p*_1_, *p*_3_). For GRN inference from singlecell protein measurements, this implies that true positive and false positives will always be observed together. Again, the kinetic conditions which allow inference of the network also confound inference via the false positive edges.

The dependence of cor(*p*_2_, *p*_3_) on *B*_1_ (cyan curves), *B*_2_ (dark-green curves) and *τ* based on (29) is shown in Fig. 3c. We have also shown the respective curves for cor(*p*_1_, *p*_2_) based on (20) there for the sake of comparison. We also performed exact stochastic simulation to verify these results as shown in Figs. 3e.

## IV. Loss of Information-Theoretic Measures for Simple GRNS

We also study the behavior of MI and the phi-mixing coefficient as a function of mRNA bursting and *τ* for the single gene and the two-gene cascade topologies using stochastic simulations ([29]). For a single-gene, like correlation, we observe a *τ*-mediated loss in MI between *m*_1_ and *p*_1_ (Fig. 4, top). For the two-gene cascade, there is *τ*-mediated loss in MI between *m*_1_ and *m*_2_ as well (Fig. 4, middle). Similar to correlation, MI between *m*_1_ and *p*_1_ appears to be larger in magnitude compared to *m*_1_ and *m*_2_. Finally, MI between *p*_1_ and *p*_2_ for the two-gene cascade in (10) exhibits an opposite trend to MI between *m*_1_ and *m*_2_, and has a *τ*-mediated loss when *τ* decreases rather than increasing (Fig. 4, bottom). The phi-mixing coefficient exhibited a similar behavior (Fig. 4b).

**Fig. 4:**
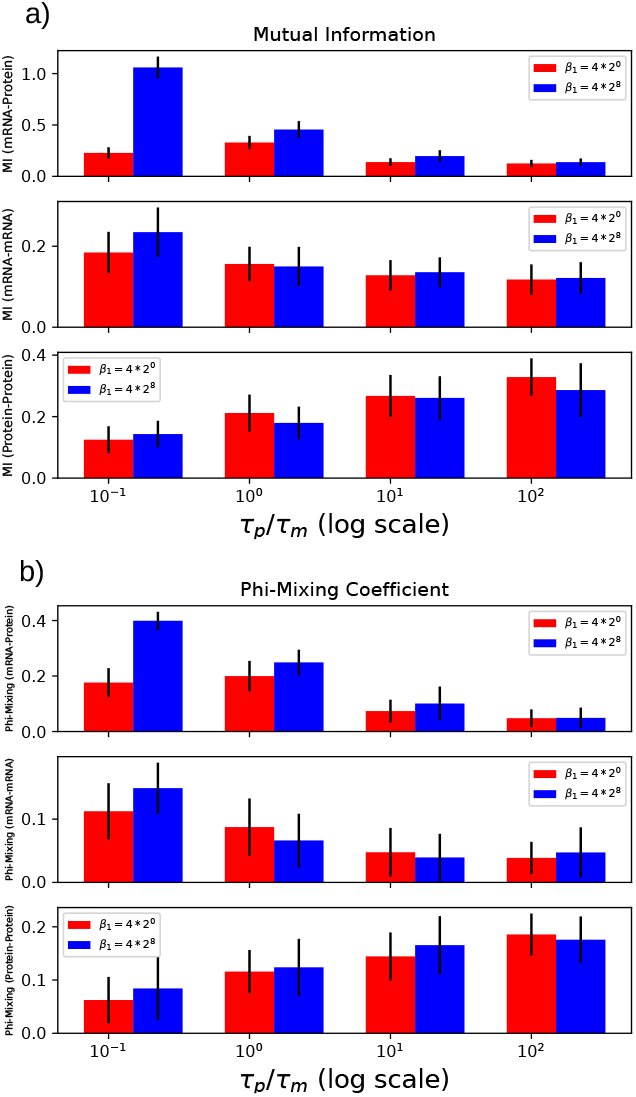
**Plot of mutual information and phi-mixing coefficient for a single-gene and the two-gene cascade** as functions of *B* (*B*_1_), and *τ*. (a) and (b) show results for mutual information (MI) and phi-mixing coefficient, respectively. (top) MI/phi-mixing coefficient between *m*_1_ and *p*_1_ for the single gene in (1) computed using exact stochastic simulations [29] for two values of *B*–one low ∝ ⟨*β*⟩= 4 (red), and one high ∝ ⟨*β*⟩ = 1024 (blue), and four different values of *τ*. (middle) MI/phi-mixing coefficient between *m*_1_ and *m*_2_ for the two-gene cascade in (10). (bottom) MI/phi-mixing coefficient between *p*_1_ and *p*_2_ for the two-gene cascade in (10). The values of other kinetic parameters are the same as before (see caption of Fig. 1). For details on the regulation function *h* (*p*_1_), see caption of Fig. 2. Error bars show one standard deviation.

## V. Loss of Inference Accuracy for real Yeast GRN in Single-Cell RNA Sequencing Data

We collected the experimentally inferred GRN for *S. cerevisiae* from the yeastract database [43]. We collected transcriptome- and proteome-wide mRNA and protein degradation rates from [44] and [45], respectively. We used scRNA-seq data generated in [46]. We only retained cells grown under complete medium conditions as degradation rate measurements were made under these conditions [44], [45]. Further, the scRNA-seq data has 12 different genotypes, including the wildtype, and we used all the genotypes.

From the GRN, we extracted edges between master regulators (TFs without any incoming regulation), and their target genes. For these edges, we computed correlation between the mRNA counts of the TF and its target. These values are shown in red in Fig. 5a. We find that these true positive edges do not violate the upper bound on such correlations. Since the TF and the target genes can have different lifetimes, we recompute the upper bound in (4), which becomes

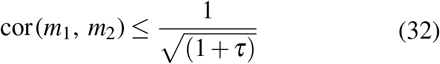

**Fig. 5:**
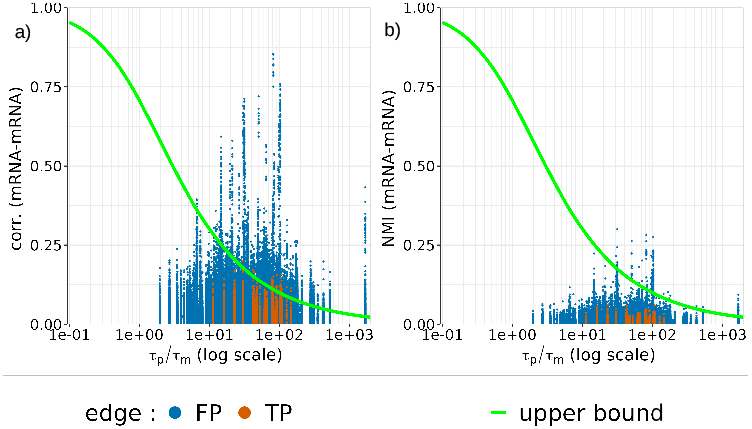
Loss in correlation and mutual information for yeast in single-cell RNA sequencing data. (a) Comparision of correlation and normalized mutual information (NMI) between a TF and its target gene (red spheres) against correlation and normalized mutual information (NMI) between two genes with common regulators but no edge between them (blue spheres) are shown in (a) and (b), respectively. The green curves represent the upper bound on correlation between a TF and its target gene given in (32)

Interestingly, allowing different lifetimes, enables the upper bound to reach the maximum possible value of 1 when *M*_1_ is much more stable than *M*_2_. The upper bound in (32) is shown by the green curve in Figs. (5)a and (5)b.

Within the limit of statistical variability, the true positive correlations from yeast scRNA-seq data do not violate (32), and consequently face a *τ*-mediated loss (Fig. (5)a). The false positive edges (blue spheres in Fig. (5)a) are not constrained by the upper bound. For false positive edges, we calculate correlation between genes which do not regulate each other, but are regulated by the master regulators. This observation shows that the insights generated from small network topologies are valid for larger GRNs as well. Further, we observed a similar behavior when we calculated normalized mutual information [47] instead of correlation for the true positive and false positive edges in Fig. (5)b.

## Discussion

We have established fundamental limits on inferrability of GRN topology from static single-cell mRNA and protein measurements. We find a relative stability-mediated loss in correlation and information-theoretic measures for mRNA species; when protein is more stable than mRNA, steady-state mRNA counts are not enough to infer the underlying GRN. This is exacerbated by the robustness of false positive correlations to this loss. The kinetic conditions which allow discovery of true positive correlations between a TF and its target gene also hinder GRN inference by amplification of false positive correlations.

We also found these constraints to be true for scRNA-seq data for yeast, *S. cerevisiae*, suggesting that the relative stability issue is true for real systems. This raises an important question on the limits of identifiability from static data. What about the dependence of other inference tasks, such as kinetic estimation, trajectory inference, clustering and differential expression, on relative stability of mRNA and protein and its propagation over GRN?

We used a simple model of gene expression, which does not incorporate complex processes such as post-transcriptional and post-transcriptional modifications. A future research direction is the establishment of constraints on GRN inferrability in more general settings.

scRNA-seq is not a static snapshot. Cells can be present in multiples states, and not not steady-state prior to sequencing. Can this be leveraged to circumvent the issues we have raised? This is an interesting problem to unpack.

GRN is essential for cellular functioning [2]–[4]. Consequently, a knowledge of its topology is important for understanding and controlling cellular functions. Experimental and computational efforts have been spent over the last two decades to unravel GRN topology for different species. However, the computational problem still remains unsolved. We have provided one explanation for this difficulty. This will motivate development of methods to circumvent the limitations we have unraveled.

